# Differential regulation of prelimbic and thalamic transmission to the basolateral amygdala by acetylcholine receptors

**DOI:** 10.1101/2021.12.28.474396

**Authors:** Sarah C. Tryon, Joshua X. Bratsch-Prince, James W. Warren, Grace C. Jones, Alexander J. McDonald, David D. Mott

## Abstract

The amygdalar anterior basolateral nucleus (BLa) plays a vital role in emotional behaviors. This region receives dense cholinergic projections from basal forebrain which are critical in regulating neuronal activity and synaptic transmission. Cholinergic signaling in BLa is thought to occur through both a slow mode of volume transmission as well as a rapid, phasic mode. However, the relative effect of each mode of signaling in BLa is not understood. Here, we used electrophysiology and optogenetics in mouse brain slices to compare regulation of afferent input from cortex and thalamus to the BLa by these two modes of transmission. Phasic ACh release evoked by single pulse stimulation of cholinergic terminals had a biphasic effect on glutamatergic transmission at cortical input, producing rapid nicotinic receptor-mediated facilitation followed by slower muscarinic receptor (mAChR)-mediated depression. In contrast, tonic elevation of ACh through application of the cholinesterase inhibitor physostigmine suppressed glutamatergic transmission at cortical inputs through mAChRs only. This suppression was not observed at thalamic inputs to BLa. In agreement with this pathway-specificity, the mAChR agonist, muscarine more potently suppressed transmission at inputs from prelimbic cortex (PL) than thalamus. Muscarinic inhibition at PL input was dependent on presynaptic M4 mAChRs, while at thalamic input it depended upon M3 mAChR-mediated stimulation of retrograde endocannabinoid signaling. Muscarinic inhibition at both pathways was frequency-dependent, allowing only high frequency activity to pass. These findings demonstrate complex cholinergic regulation of afferent input to BLa that depends upon the mode of ACh release and is both pathway specific and frequency dependent.

**Significance statement:** Cholinergic modulation of the basolateral amygdala regulates formation of emotional memories, but the mechanisms underlying this regulation are not well understood. Here, we show, using mouse brain slices, that ACh differentially regulates afferent transmission to the BLa depending on the mode of cholinergic signaling. Rapid, phasic ACh produces a biphasic excitatory-inhibitory regulation of glutamatergic transmission mediated by nicotinic and muscarinic receptors, respectively. In contrast, slow, tonic ACh produces muscarinic inhibition only. Tonic regulation is pathway specific with cortical input regulated more strongly than thalamic input. This disparity is caused by differential regulation by M4 and M3 receptors at the two inputs. Specific targeting of these receptors may thus provide a therapeutic strategy to bias amygdalar processing and regulate emotional memory.

## Introduction

The basolateral amygdala is a brain region central to emotional processing and is necessary for associating cues with both positive and negative valence outcomes (LeDoux et al., 1990; Baxter and Murray, 2002; Janak and Tye, 2015). Compared to other brain regions, the basolateral amygdala, especially the anterior subdivision of the basolateral nucleus (BLa), receives the densest cholinergic projections from basal forebrain (BF; Woolf, 1991; Muller et al., 2011; Zaborszky et al., 2012), suggesting that acetylcholine (ACh) plays a central role in regulating neurons in this region. Indeed, cholinergic mechanisms in the BLa are important modulators of emotional memory (Power et al., 2003; McGaugh, 2004; Jiang et al., 2016; Wilson and Fadel, 2017). Cholinergic mechanisms in the BLa are also thought to regulate reward devaluation learning (Salinas et al., 1997), performance in tests of anxiety and depression-like behaviors (Mineur et al., 2016, 2018), and conditioned cue reinstatement of cocaine seeking (See et al., 2003; See, 2005), suggesting roles for ACh in the BLa in anxiety and fear disorders and drug addiction. These findings underscore the impact of cholinergic activity in the BLa in the pathophysiology of several neuropsychiatric diseases and highlight the need to better understand cholinergic modulation of the BLa.

ACh is thought to modulate neuronal circuits through both a slow mode of volume transmission as well as a more temporally precise phasic mode and thereby regulate neural activity over a range of temporal and spatial scales (Disney and Higley, 2020; Sarter and Lustig, 2020). These two modes of cholinergic transmission likely engage different types of ACh receptors with different kinetics, affinities, desensitization characteristics and cellular locations. Thus, the nature of the cholinergic response may depend upon the dynamics of ACh release. In the BLa cholinergic signaling shapes neural activity through multiple mechanisms, including the regulation of presynaptic release probability (Jiang and Role, 2008; Jiang et al., 2016). Cholinergic projections from the BF converge with excitatory terminals providing an anatomical basis for cholinergic regulation of glutamatergic transmission in this area (Muller et al., 2011, 2013). As in other brain regions, ACh in the BLa is thought to act on presynaptic nicotinic receptors to enhance glutamate release (Jiang and Role, 2008; Jiang et al., 2016) and on muscarinic receptors to suppress release (Sugita et al., 1991; Yajeya et al., 2000). However, prior studies examining cholinergic modulation have used exogenous agonists or sustained optogenetic stimulation of cholinergic afferents, which results in broad spatial and temporal activation of cholinergic receptors. Evidence that cholinergic regulation can occur in a rapid and precise timescale sufficient to modulate individual synaptic events is lacking. This is significant as cholinergic neurons in BF can exhibit fast and precise responses to behaviorally relevant cues (Hangya et al., 2015; Crouse et al., 2020). Furthermore, little is known about the types of cholinergic receptors engaged by different modes of release or the relative role of ACh in modulating different afferent inputs to this region.

In the present study we have investigated cholinergic regulation of afferent input to the BLa in mouse brain slices. The BLa receives major excitatory projections from prelimbic cortex (PL) and midline thalamic nuclei (MTN) which are thought to play distinct roles in amygdalar-dependent behaviors (Corcoran and Quirk, 2007; Arruda-Carvalho and Clem, 2014; Salay et al., 2018; Amir et al., 2019; Ahmed et al., 2021). We find that endogenously released ACh from single pulse optical stimulation can rapidly and precisely regulate glutamatergic transmission at cortical inputs, suggesting that cholinergic neuromodulation can serve precise, computational roles in the BLa at this timescale. This modulation differs from that during sustained ACh exposure, indicating involvement of different ACh receptors. During tonic ACh exposure, cholinergic regulation is pathway-specific, producing stronger regulation of cortical than thalamic input. It is also frequency dependent and acts as a high pass filter for incoming signals. Through these mechanisms, ACh dynamically shapes afferent input to BLa to bias amygdalar processing of salient cues.

## Materials and Methods

### Animals

All experiments were performed in adult ChAT-Cre mice (B6;129S6-Chat^tm2(cre)Lowl/J^; JAX Stock No. #006410) of either sex. These mice express Cre-recombinase under the control of the choline acetyltransferase gene. Alternately, in some experiments the adult F1 progeny of ChAT-Cre mice crossed with Ai32 mice (B6;129S-Gt(ROSA)26SOR^tm21(CAG-COP4*H134R/EYFP)Hze^/J, JAX Stock No. #012569) were used. Mice were group housed in a climate-controlled facility with a 12/12 light/dark cycle and provided with *ad libitum* access to food and water. All animal care and use procedures were approved by the University of South Carolina’s Institutional Animal Care and Use Committee and performed in compliance with the guidelines approved by the National Institution of Health Guide for the Care and Use of Laboratory Animals (Department of Health and Human Services).

### AAV delivery

Mice 1.5-3 months old were anesthetized under deep isoflurane anesthesia and placed in a stereotaxic surgery device (Stoelting, Wood Dale, IL). For *ex-vivo* slice electrophysiology experiments utilizing released ACh, 0.2μL of rAAV5-EF1a-DIO-hChR2(H134R)-EYFP (UNC Viral Vector Core, Chapel Hill, NC) was delivered bilaterally into the BF, including the ventral pallidum/substantia innominata (VP/SI; from Bregma: AP 1.2 mm; ML +1.3 mm; DV −5.3mm), the main source of cholinergic inputs to the BLa. For *ex-vivo* slice electrophysiology experiments examining PL input to the BLa, 0.15μL of rAAV5-CAMKII-hChR2(H134R)-EYFP-WPRE (UNC Viral Vector Core, Chapel Hill, NC) was delivered bilaterally to the PL (from Bregma: AP 1.9 mm; ML +0.3 mm; DV −2.0 mm). For experiments examining MTN input to the BLa, single injections of 0.2μL of rAAV5-CAMKII-hChR2(H134R)-EYFP-WPRE (UNC Viral Vector Core, Chapel Hill, NC) were delivered to the MTN (from Bregma: AP −0.3 mm; ML 0.0 mm; DV −3.9 mm). For experiments examining ventral subicular (vSUB) input to the BLa, single injections of 0.2μL of rAAV5-CAMKII-hChR2(H134R)-EYFP-WPRE (UNC Viral Vector Core, Chapel Hill, NC) were delivered to the vSUB (from Bregma: AP −2.5 mm; ML +3.2 mm; DV −5.3 mm). Mice were used for experiments at least 3 weeks after surgery.

### Immunofluorescence

To validate expression of channelrhodopsin in BF cholinergic neurons, ChAT-Cre mice injected in the BF with rAAV5-EF1a-DIO-hChR2(H134R)-EYFP or ChAT-Cre/Ai32 mice were transcardially perfused with ice cold phosphate buffered saline containing 0.5% nitrite followed with 4% paraformaldehyde. Brains were post-fixed overnight in 4% paraformaldehyde at 4°C. 50 μm coronal brain sections were cut using a vibratome (VT1200S, Leica, Nussloch, Germany). Slices were blocked in TBS containing 0.5% Triton X-100 and 10% normal donkey serum and incubated for 30 minutes at room temperature. Sections were then incubated for 48 hours at room temperature in goat anti-ChAT primary antibody (1:1000, AB144P, Millipore). Following rinse, sections were again incubated at room temperature for 3 hours in TBS containing an Alexa Fluor 546-conjugated donkey anti-goat IgG secondary antibody (1:400, A-11056, ThermoFisher), 0.5% Triton X-100, and 2% normal donkey serum. Sections were rinsed and mounted on slides with ProLong diamond antifade mountant DAPI (ThermoFisher, P36971) and imaged on a Leica SP8 Multiphoton confocal microscope (Leica Microsystems). ChR2-EYFP was assessed by the endogenous EYFP fluorescent signal. The number of neurons positive for both EYFP and ChAT, or EYFP alone, were counted in each image using ImageJ (NIH).

### Slice preparation

Mice were deeply anesthetized with isoflurane and the brain quickly extracted and submerged in ice-cold (4°C) ‘cutting’ artificial cerebrospinal fluid (aCSF) containing (in mM): 110 choline chloride, 2.5 KCL, 25 NaHCO_2_, 1.0 NaH_2_PO_4_, 20 glucose, 5 MgCl_2_, 0.5 CaCl_2_, 1-5 kynurenic acid and continuously bubbled with 95% O_2_/5% CO_2_. Brains were cut into 300μm thick (for whole-cell experiments) or 500μm thick (for field recordings) coronal sections using a vibratome (VT1000S, Leica, Nussloch, Germany). Slices were transferred to an incubation chamber filled with warmed ACSF containing (in mM): 125 NaCl, 2.7 KCl, 25 NaHCO_2_, 1.25 NaH_2_PO_4_, 10 glucose, 5 MgCl_2_, 0.5 CaCl_2_ and bubbled with 95% O_2_/5% CO_2_ at 34-36°C. In some experiments 1-5 kynurenic acid was included in the incubating ACSF. After a minimum of 20 minutes, the incubation temperature was allowed to equilibrate to room temperature for at least 40 min before use in recording.

### Slice electrophysiology recordings

For recording, slices were placed in a submersion chamber and continuously perfused with oxygenated (95% O_2_/5% CO_2_), recording ACSF a rate of 4-6 mL/min (for field potentials) or 1-2 mL/min (for whole cell) at 30-32°C. Recording ACSF contained (in mM): 125 NaCl, 2.7 KCl, 25 NaHCO_2_, 1.25 NaH_2_PO_4_, 10 glucose, 1 MgCl_2_, 2 CaCl_2_ (pH 7.4, 305 mOsm). Field potentials were recorded from the BLa with an Axoprobe 1A amplifier (Molecular Devices, Sunnyvale, CA) using borosilicate glass electrodes which had a resistance of 1-3 MΩ when filled with recording ACSF. For whole cell recording, pyramidal neurons in the BLa were visualized using infrared-differential interference contrast optics through a 40x objective (Olympus BX51WI). Borosilicate glass electrodes of 4-6 MΩ resistance were used for recordings and filled with a potassium gluconate internal solution consisting of (in mM): 135 K-gluconate, 5 KCl, 10 HEPES, 2 MgCl_2_, 2 MgATP, 0.3 NaGTP, 0.5 EGTA. Voltage clamp recordings were made at a holding potential of - 70 mV with a Multiclamp 700B (Molecular Devices, Sunnyvale, Ca) amplifier. Experiments were discarded if significant changes occurred in input or series resistance which were monitored throughout. All responses were filtered at 1kHz, digitized using a Digidata 1440A A-D board (Molecular Devices, Sunnyvale, CA) and analyzed using pClamp10 software (Molecular Devices, Sunnyvale, Ca).

To evoke glutamatergic field EPSPs (fEPSPs) PL or MTN fibers expressing hChR2(H134R)-EYFP were stimulated with single or dual (50 ms apart) rectangular pulses (1-3 ms duration) of 490nm blue LED light (M490F3, ThorLabs Inc, Newton, New Jersey) delivered through a fiber optic cable directly over the recording site in the BLa every 30 seconds. For whole cell recording the pulse of 470 nm blue light (pE-4000, CoolLED, Andover, UK) was delivered through the 40x objective. Alternately, in some experiments fEPSPs or whole cell EPSCs in BLa were electrically evoked using a 0.1ms rectangular current pulse delivered through a monopolar platinum-iridium stimulating electrode (FHC Inc, Bowdoin, ME) placed in the external capsule (EC). Glutamatergic responses were pharmacologically isolated by blocking GABA_A_ receptors (10μM-100μM picrotoxin or 10μM bicuculline), GABA_B_ receptors (2μM CGP55845) and N-methyl-D-aspartate (NMDA) receptors (50μM L-2-amino-5-phosphonovaleric acid (D-APV) or 10μM MK801). 6-cyano-7-nitroquinoxaline-2,3-dione (CNQX, 25μM) was added at the conclusion of some experiments to confirm that the response was mediated by glutamate receptors.

To study the effects of released ACh, cholinergic fibers expressing hChR2(H134R)-EYFP were optogenetically stimulated with 2-3ms pulses of 490nm blue LED light delivered directly over the recording site in the BLa every 90 seconds. A single light pulse or a theta burst of light [4 bursts of light (4 pulses at 50 Hz) delivered every 200 ms] was delivered either immediately before or 250 ms before electrical stimulation. In whole cell experiments to determine the effect of light on the EPSC amplitude, direct post-synaptic currents produced by optically released ACh alone were recorded and subtracted from evoked EPSC traces where light was applied.

#### Drugs

Baclofen, muscarine chloride, N-ethylmaleimide, and physostigmine were purchased from Millipore Sigma (St. Louis, MO). Bicuculline, D-AP5, CNQX, CGP55845 hydrochloride, MK 801 maleate, mecamylamine hydrochloride and AM251 were purchased from HelloBio (Princeton, NJ). WIN 55,212-2, oxotremorine M, and VU10010 were purchased from Tocris Biosciences (Bristol, UK). 4DAMP, AF-DX 116, VU0255025, and Atropine were purchased from Abcam (Cambridge, UK). AM630 was purchased from Cayman Chemical Company (Ann Arbor, MI) and VU0467154 was purchased from StressMarq Biosciences (Victoria, BC). All reagents were added from freshly prepared stock solution to the ACSF. Drugs were applied using bath perfusion and drug concentration in the bath during wash-in was allowed to equilibrate before measurements were taken.

### Data Analysis and Statistics

Electrophysiological data analysis was performed using pClamp 10 (Molecular Devices) and OriginPro 2018b (Microcal, Northampton, MA) software. For released ACh experiments, consecutive sweeps in “Light ON” or “Light OFF” conditions (2-6 sweeps) were averaged and the peak amplitude of the averaged EPSC or fEPSP was measured. For experiments involving optogenetically stimulated PL and MTN input and bath application of muscarine, the peak amplitude of fEPSPs was measured as the average peak amplitude of the steady-state evoked fEPSP response in each pharmacological condition. All peak amplitudes were normalized to the baseline condition (“control”) and are expressed as the mean ± SEMs. Concentration–response curves represent a least-squares fit of each data set to a sigmoidal (logistic) curve (Graph-Pad Prism, GraphPad Software, San Diego, California). The IC_50_ and Hill slope were calculated from this curve. Means, standard errors and 95% confidence intervals (95% c.i.) were determined by the fitting algorithm. In some experiments, multiple slices per animal were used, so for all experiments n=slice number and N=animal number. Statistical significance was determined using a Student’s *t* test (paired or unpaired) or a one-way analysis of variance (ANOVA) with a *post-hoc* Tukey test (α < 0.05 was taken as significant).

## Results

### Immunofluorescent verification of ChR2 Expression in BLa-projecting cholinergic neurons in the BF

In order to optogenetically evoke ACh release, two strategies were used to selectively express channelrhodopsin in BF cholinergic axons in BLa. First, ChR2-EYFP was expressed in BF cholinergic neurons of ChAT-Cre mice through Cre-dependent rAAV-mediated transfection (Unal et al., 2015; Aitta-aho et al., 2018). Four weeks after AAV injection, we verified selective ChR2-EYFP expression in neurons labelled with ChAT antibody (ChAT+) in the BF (Fig. 1). Cell counts of ChAT+ neurons, ChR2-EYFP+ neurons (ChR2+), or neurons expressing both ChAT+ and ChR2-EYFP+ at the injection site revealed that most ChAT+ neurons expressed ChR2-EYFP (70.2 ± 4.26%, N = 5, Figure 1). Furthermore, immunoreactivity for ChAT in the majority of ChR2-EYFP+ cells (89.8 ± 2.4%, N = 5) confirmed that expression of ChR2 was restricted to cholinergic neurons in this region (Fig. 1B, bottom). Axons from labelled ChAT+ neurons in BF densely innervated the BLa (Fig. 1A), as previously described (Aitta-aho et al., 2018), further supporting the selective labelling of cholinergic projections to BLa.

**Figure 1.**
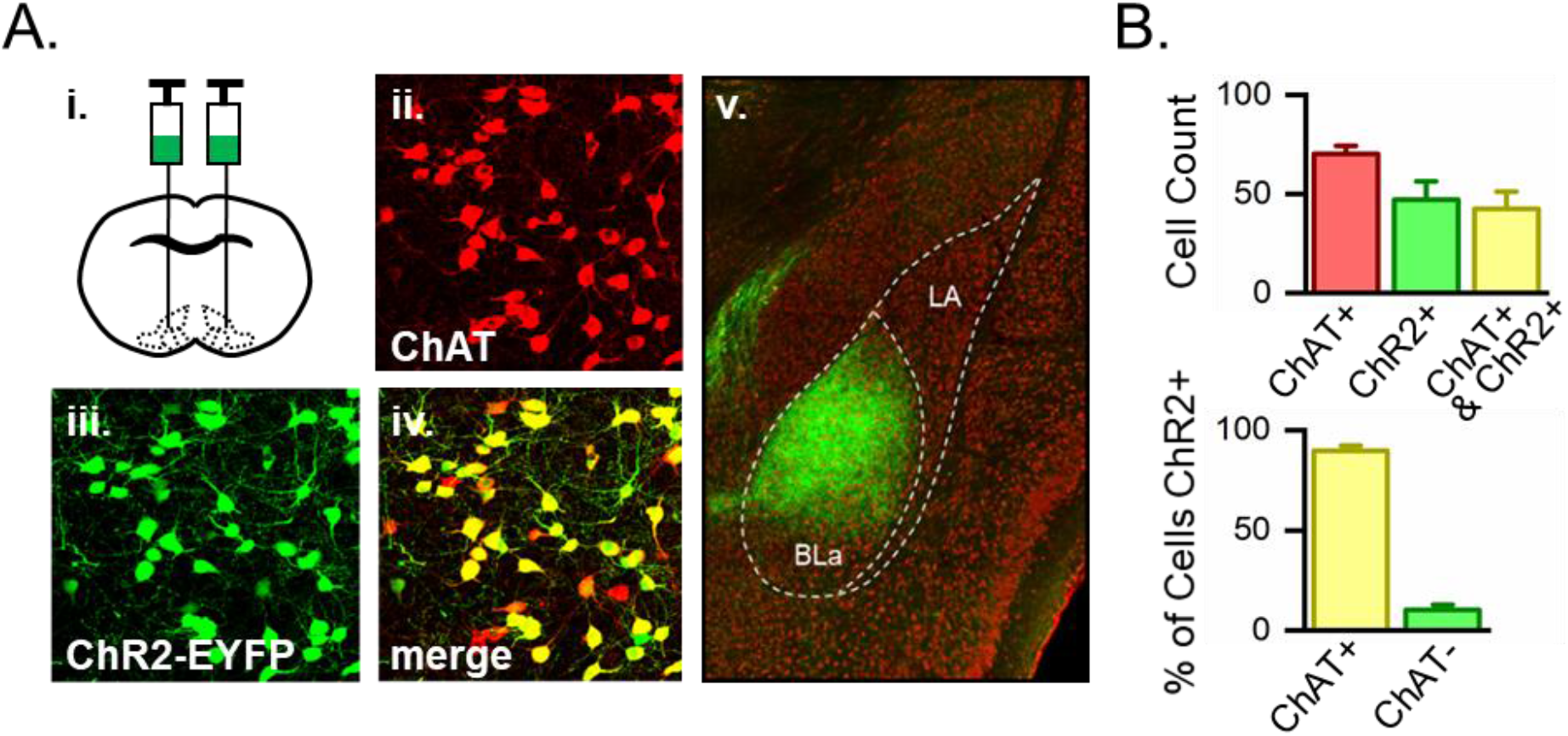
ChR2 expression in ChAT+ neurons. **A.** Viral injection into the BF of ChAT-cre mice led to ChR2-EYFP expression in ChAT-containing neurons that project to BLa. i. schematic of injection sites, ii. ChAT-immunopositive cell bodies (red) at the injection site, iii. EYFP-labeled ChR2+ cells (green) at the injection site, iv. Merged image showing ChR2-EYFP cells are immunopositive for ChAT (yellow). v. In the same mouse ChR2-EYFP-expressing axons (green) strongly innervate the BLa. **B.** (Top) Counts of BF neurons per 50μm thick coronal tissue section that were labeled with either ChAT+ (red), ChR2-EYFP+ (green) or both (yellow). (Bottom) The majority of cells expressing ChR2-EYFP (89.8 ± 2.4%, N = 5) were also immunopositive for ChAT.

Channelrhodopsin was also expressed in ChAT+ neurons using a double transgenic strategy in which ChAT-Cre mice were crossed with an Ai-32 reporter mouse line expressing Cre-dependent ChR2-EYFP (Hedrick et al., 2016; Baker et al., 2018). In the F1 generation of these mice (ChAT-Cre/Ai32 mice) the majority of BF ChAT+ neurons (80 + 3%, 877 cells, N = 3) were immunopositive for ChR2-EYFP. Furthermore, immunoreactivity for ChAT in most ChR2-EYFP-immunopositive cells (99.1 ± 0.8%, 694 cells, N = 3) confirmed that expression of ChR2 was restricted to cholinergic neurons. Notably, the BLa of these mice did not contain any cell bodies positive for ChR2, ensuring selective activation of BF derived cholinergic terminals with optogenetic stimulation during BLa slice recordings.

### Synaptically released acetylcholine biphasically regulates cortico-amygdalar transmission in the BLa through both nicotinic and muscarinic receptors

In vivo recordings indicate that a behaviorally relevant cue can recruit BF cholinergic neurons to synchronously fire a single, precisely time spike or brief burst of action potentials (Hangya et al., 2015). To determine the effect of this cholinergic neuron activity on afferent input to the BLa, cholinergic terminals were stimulated with a single blue light pulse (470 nm) and the effect on synaptic transmission at cortical inputs to BLa in ChAT-Cre/Ai32 mice examined. EPSCs were evoked in BLa pyramidal cells by stimulation of cortical afferents in the external capsule (EC, Fig 2A; Jiang et al., 2016). Optogenetic stimulation of cholinergic terminals had a biphasic effect on the amplitude of this EPSC in the majority of cells (Fig. 2). Stimulation of cholinergic terminals immediately before stimulation of cortical afferents (Early interval) evoked a facilitation of the EPSC. This early facilitation was sensitive to the frequency at which cholinergic terminals were stimulated. In these experiments cholinergic terminals were stimulated at a frequency of 0.011 Hz, as higher frequency stimulation resulted in a rundown or loss of the facilitation. The extent of the facilitation varied between cells (range: 92-137%, mean: 107.5 + 2%, n=23, N=10) with 17 of 23 cells (73.9%) exhibiting a facilitation (Fig 2C). Increasing the interval between the light pulse and cortical afferent stimulation caused this facilitation to rapidly diminish and become a depression at intervals greater than 20 ms. When the cortical afferents were stimulated 250 ms after the light pulse, the EPSC was inhibited (range: 46-96%; mean: 79.4 ± 3%; n=17, N = 10; Fig 2C-D). All cells that exhibited early facilitation also exhibited late inhibition. However, EPSCs in five cells exhibited late inhibition with no early facilitation. Late inhibition was similar in amplitude whether cholinergic terminals were stimulated with a single light pulse or a theta burst (4 pulses at 50 Hz) of light pulses (Fig 2C).

**Figure 2.**
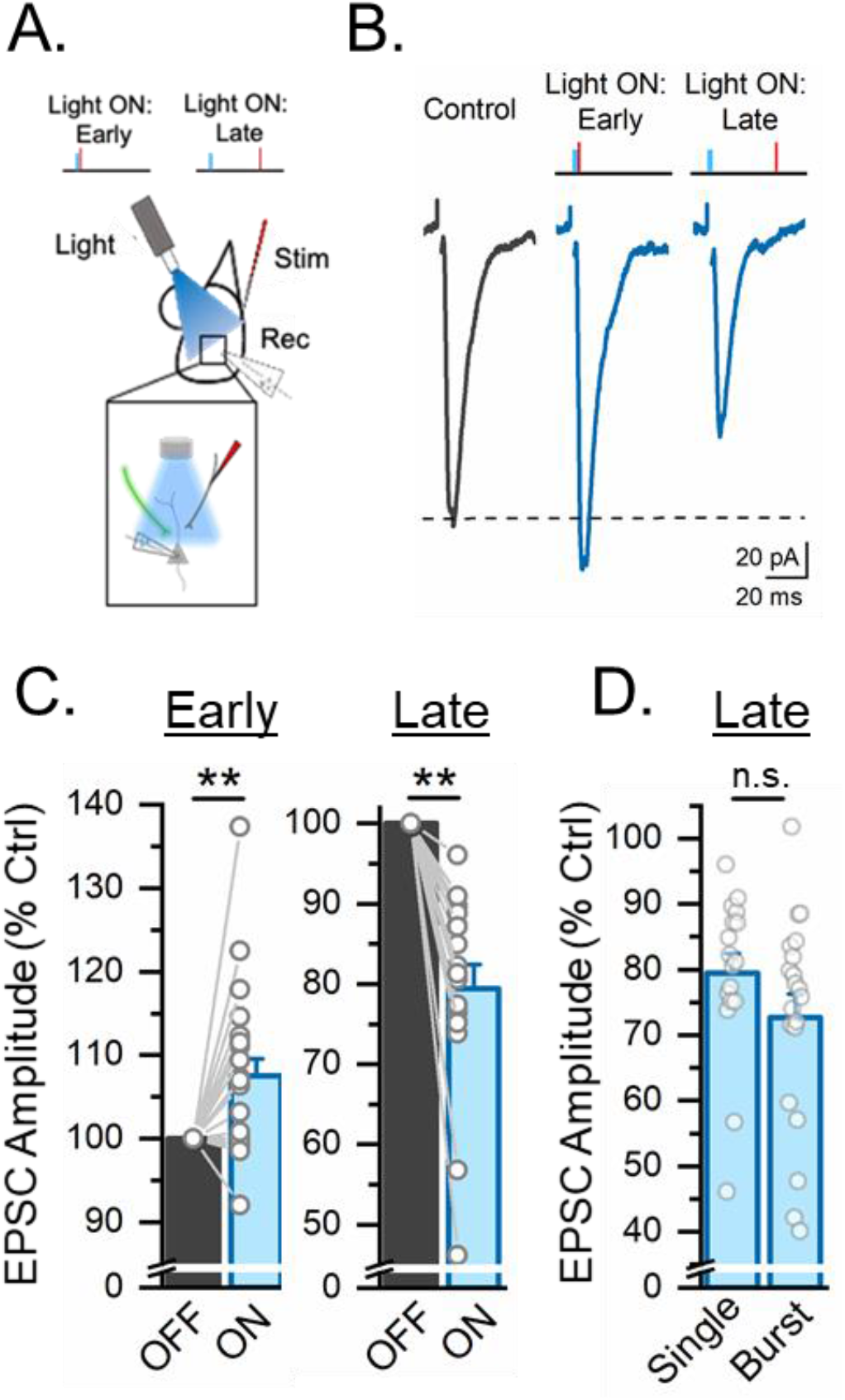
Released ACh exerts a biphasic effect on the cortical EPSC in BLa. **A.** Schematic illustrating the placement of a stimulating electrode in the external capsule and recording electrode in the BLa. Blue light pulses (470nm, 1 ms) were delivered above the recording site to stimulate cholinergic terminals prior to external capsule (EC) stimulation. (Top) “Light ON: Early” illustrates an EC stimulus delivered immediately after a single light pulse. “Light ON: Late” illustrates an EC stimulus delivered 250 ms after a single light pulse. **B,C.** Optogenetic activation of cholinergic terminals evoked facilitation of the EC-evoked EPSC at the early interval (n=23, N=10; paired t-test, p<0.01) and inhibition at the late interval (n=17, N=10; paired t-test, p<0.01). The extent of facilitation or inhibition varied between cells (open circles). Facilitation at the early interval was absent in 6 cells, while inhibition was present at the late interval in all cells. **D.** Inhibition at the late interval was similar (two sample t-test, p = 0.17) whether it was evoked by a single pulse (1 ms, n = 17) or burst of blue light pulses (4 bursts of light (4 pulses at 50 Hz) delivered every 200 ms, n = 20). **p<0.01, n.s. not significant.

Pharmacological analysis revealed that the early facilitation by ACh was completely blocked by the nicotinic antagonist, mecamylamine (10 μM; Fig. 3A), indicating that it was nAChR-mediated. Mecamylamine had no effect on the EPSC amplitude at the 250 ms interval (Fig 3C), demonstrating the absence of any delayed effect of nAChRs on the EPSC, as has been reported in cortex (Urban-Ciecko et al., 2018). Nicotinic receptors are prone to desensitization during sustained increases in extracellular ACh or in response to prolonged application of agonists (Giniatullin et al., 2005). Thus, the observed lability of the early facilitation is consistent with nicotinic receptor desensitization limiting the potentiation during higher stimulus frequencies. The site of action of nAChRs was investigated by examining the effect of cholinergic stimulation on paired pulse facilitation. Nicotinic facilitation significantly reduced the paired pulse ratio at the early interval (Fig 3B), indicating a presynaptic site of action in agreement with prior studies (Jiang and Role, 2008; Cheng and Yakel, 2014; Tang et al., 2015). In contrast, late cholinergic suppression of the EPSC was blocked by bath application of the muscarinic antagonist, atropine (5 μM; Fig. 3C), demonstrating that it was mAChR-mediated. This mAChR-mediated depression of the EPSC significantly increased the paired pulse ratio at the later interval (Fig. 3D), suggesting that the mAChRs were also presynaptic.

**Figure 3.**
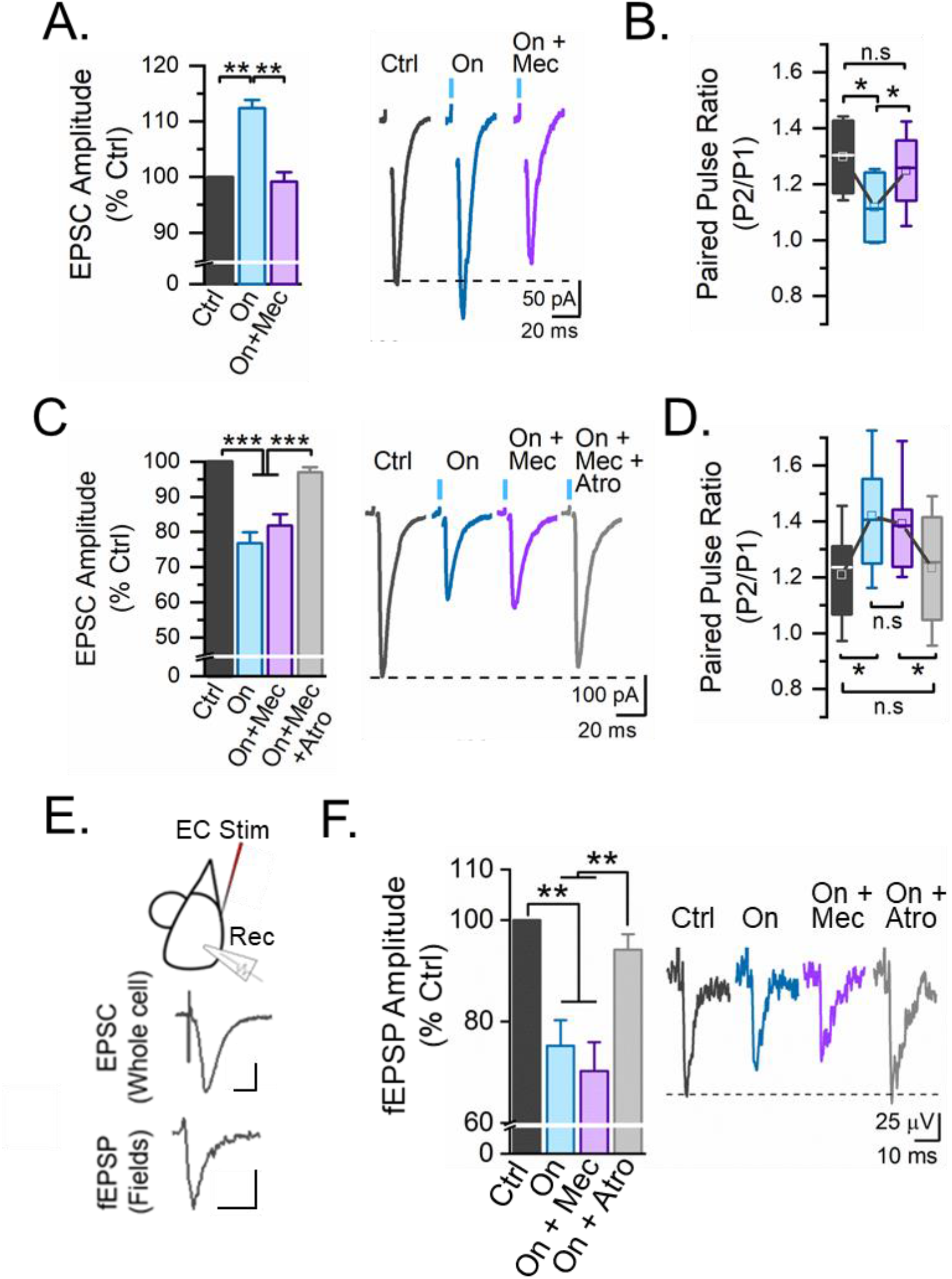
Released ACh regulates glutamatergic input to the basolateral amygdala through presynaptic nicotinic and muscarinic receptors. **A.** Mecamylamine (Mec, 10μM) blocks facilitation of the EPSC at the early interval, indicating that this facilitation is nAChR-mediated (n=12; N=8; one way ANOVA, F(2,33)=22.14; p < 0.001). **B.** At the early interval, ACh-induced reduction of the paired pulse ratio is reversed by Mec (n = 5; N = 5; paired t-test; p < 0.05). **C.** Cholinergic inhibition of the EPSC at the late interval is blocked by atropine (Atro; 5 μM), but not Mec (n = 11; N = 10; one way ANOVA, F(3,40)=28.30; p < 0.0001). **D.** At the late interval the ACh-evoked increase in the paired pulse ratio is reversed by atropine (n = 5, N = 5; student’s t-test; p < 0.05). **E.** (Top) Schematic illustrating placement of the stimulating electrode in external capsule and recording electrode in BLa. EC stimulation evoked an EPSC when recording from a BLa pyramidal neuron (Middle) or a field EPSP (fEPSP, Bottom) when recording from an extracellular field electrode. Calibration: 75 pA, 75 μV, 10 ms. **F.** Optogenetic activation of cholinergic terminals produced an atropine-sensitive inhibition of the fEPSP evoked by EC stimulation 250 ms later (n = 7; N = 7; one way ANOVA, F(3,24)=12.34; p < 0.001). *p<0.05, **p<0.01,***p<0.001, n.s. not significant.

A similar cholinergic-induced late inhibition of cortical-evoked transmission was also evident in field potential recordings in the BLa. As shown in Fig. 3E, EC stimulation evoked field EPSPs (fEPSPs) in BLa that reflected EPSCs in pyramidal neurons during whole cell recording. Optogenetic stimulation of cholinergic terminals with theta burst stimulation [4 bursts of light (4 pulses at 50 Hz) delivered every 200 ms] significantly inhibited the fEPSP evoked by EC stimulation 250 ms later. Theta burst stimulation was chosen for these studies to reflect BF activity during active waking and paradoxical sleep (Lee et al., 2005). Cholinergic inhibition of the fEPSP was unaffected by mecamylamine (10 μM), but was completely reversed by application of atropine (5 μM; Fig. 2F), indicating that it was muscarinic receptor-mediated. Taken together, these findings suggest that single pulse stimulation of ACh terminals evokes a biphasic modulation of cortical input by ACh, whereby ACh acts through a precisely timed action on presynaptic nAChRs receptors to rapidly facilitate cortical neurotransmission to the BLa and on presynaptic mAChRs to cause a delayed suppression. In contrast, during theta pattern train stimulation, nicotinic receptors desensitize, leaving only a monophasic inhibition of cortical input mediated by muscarinic receptors.

### Tonic acetylcholine differentially regulates cortical and thalamic input to the BLa

A behaviorally salient cue can recruit BF cholinergic neurons to fire, producing a phasic release of ACh into BLa (Aitta-aho et al., 2018; Crouse et al., 2020). In contrast, during prolonged emotional arousal extracellular acetylcholine levels in the amygdala exhibit a sustained increase (Kellis et al., 2020). To investigate the impact of this increased tonic acetylcholine on synaptic transmission, we increased extracellular ACh by applying physostigmine to inhibit acetylcholinesterase, the enzyme that catalyzes the breakdown of ACh. We compared the effect of increasing concentrations (0.3-10μM) of physostigmine on the amplitude of the EC-evoked fEPSP. Blocking AChE led to a concentration-dependent suppression of the EC-evoked fEPSP (Fig 4A,B). Antagonism of muscarinic receptors (5 μM atropine) reversed this suppression, indicating that the inhibition was muscarinic receptor-mediated. The ability of AChE inhibition to suppress the EC-evoked fEPSP demonstrates the presence of tonically released ACh in the brain slice and suggests that the impact of released ACh on synaptic transmission in this pathway is limited by this enzyme, in line with previous studies (Aitta-aho et al., 2018).

**Figure 4.**
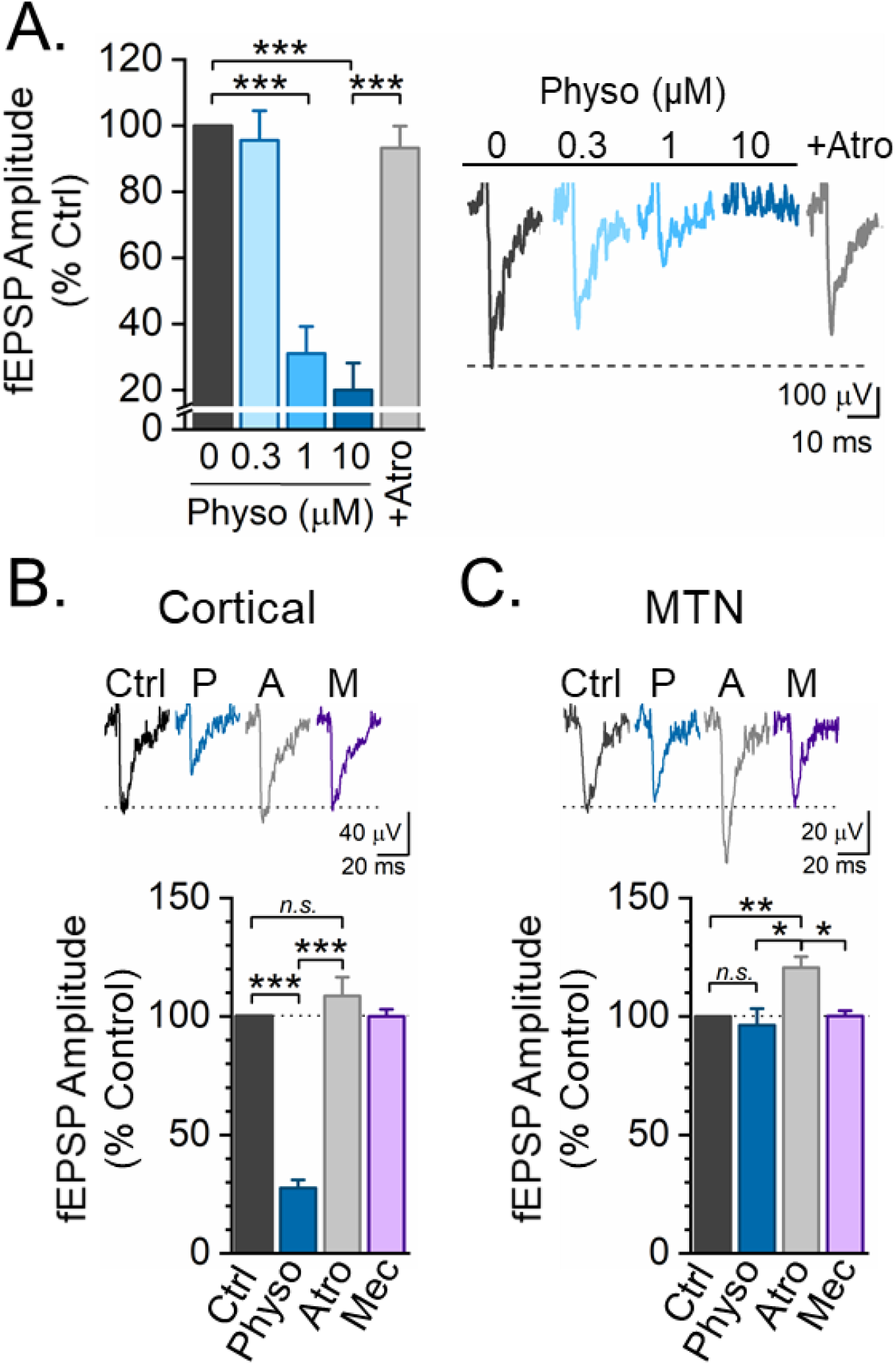
Pathway specific regulation of afferent input to BLa by tonic ACh. **A.** Physostigmine (Physo) inhibits the EC-evoked fEPSP in a concentration dependent manner. This inhibition is reversed by atropine (0.3 μM Physo: n=7, N=7, 1 μM Physo: n=5, N=5; 10 μM Physo: n=5, N=5; 5 μM atropine: n=4, N=4; one-way ANOVA, F(4,23)=28.73; p<0.0001). **B,C.** In a separate group of mice the effect of Physo (10 μM) on cortical and MTN fEPSPs recorded at the same site in BLa was compared. **B.** Physo strongly inhibited the cortical fEPSP evoked by EC stimulation. This inhibition was blocked by Atro (5 μM), but unaffected by Mec (10 μM), indicating a role for mAChRs, but not nAChRs (one-way ANOVA, F(3,17)=82.25; p<0.001). **C.** Physo had little effect on the fEPSP evoked by optogenetic stimulation of MTN input. In contrast, Atro blocked mAChRs, revealing an underlying potentiation that was subsequently inhibited by Mec (one-way ANOVA, F(3,22)=5.93; p<0.01). These results suggest that the elevation of tonic ACh produced by Physo engaged both nAChRs and mAChRs to produce opposing and offsetting effects at MTN synapses. During elevation of tonic ACh, mAChR inhibition is stronger at cortical input, whereas nAChR-mediated facilitation is stronger at MTN input. *p<0.05, **p<0.01, ***p<0.001, n.s. not significant.

**Figure 5.**
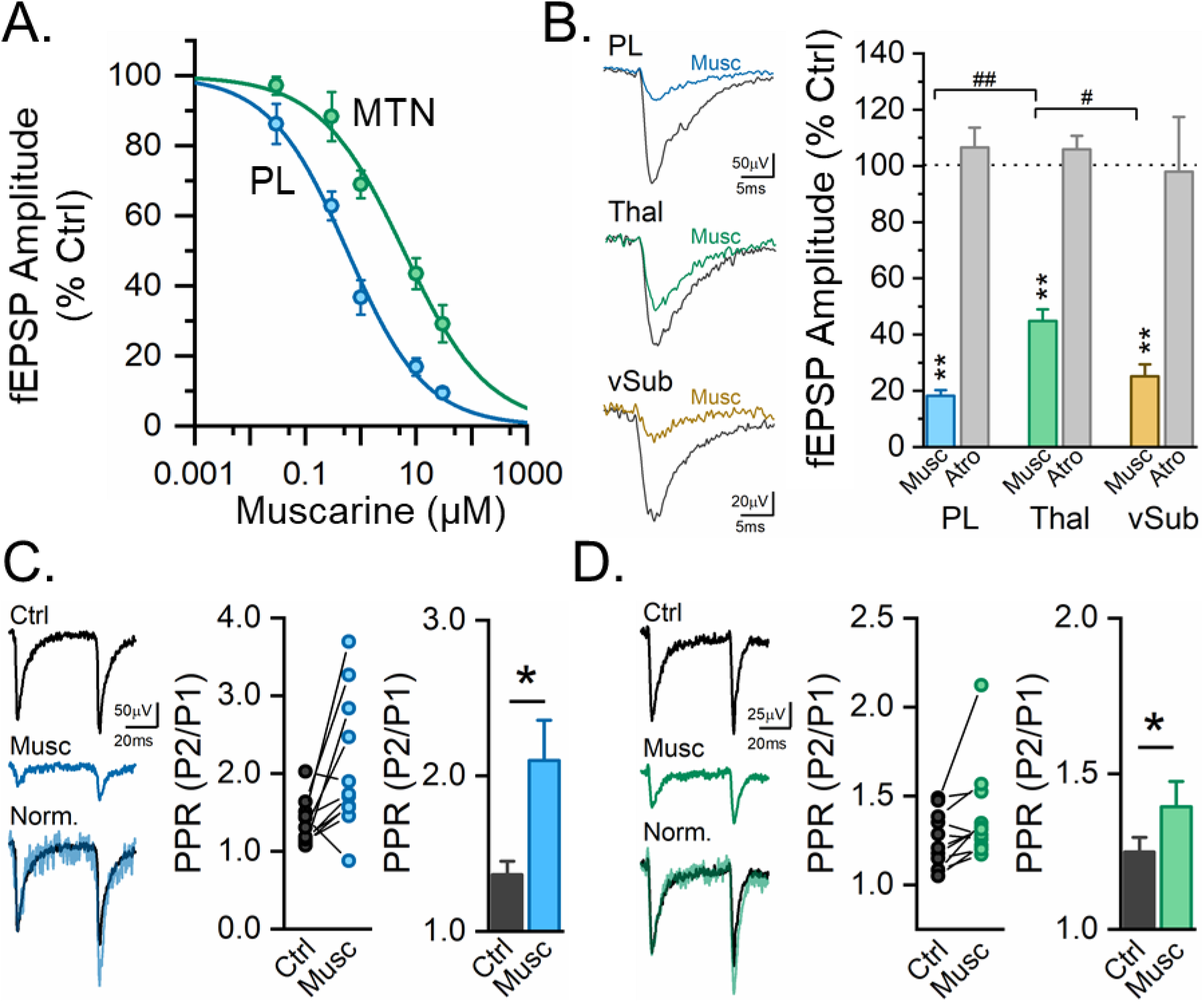
Stimulation of presynaptic muscarinic receptors more strongly inhibits cortical and subicular projections, than MTN projections to the BLa. **A.** Effect of muscarine (0.3-10 μM) on the fEPSP evoked by optogenetic stimulation of either prelimbic or MTN input to BLa. Muscarine produced a concentration dependent inhibition of the fEPSP in both pathways. The inhibitory effect of muscarine in the MTN pathway was shifted significantly to the right (PL Input, n = 5-35; MTN input, n = 4-27). **B.** Selective optogenetic stimulation of PL, MTN or vSub input to the BLa evoked a fEPSP which was inhibited by muscarine (10 μM). Muscarine produced significantly greater inhibition of PL and vSub, than MTN input (PL, n = 32, N = 27; MTN, n = 26, N = 22; vSUB, n = 5, N = 5; One way ANOVA, F(2,60)=20.10; p < 0.001). **C, D.** Muscarine inhibited the first fEPSP, but significantly enhanced paired pulse facilitation (50 ms interstimulus interval) at both the PL (PL, n=11; N=11; Students t-test, p < 0.02) and MTN inputs (n=11; N=11; Student’s t-test, p < 0.05), indicating a presynaptic site of action in each pathway. *p<0.05; **p<0.01; #p<0.05; ##p<0.01.

Inputs from both cortex and midline thalamic nuclei exert significant influence over BLa activity to regulate amygdalar responses to emotionally arousing stimuli (Corcoran and Quirk, 2007; Arruda-Carvalho and Clem, 2014; Salay et al., 2018; Amir et al., 2019; Ahmed et al., 2021). Cholinergic mechanisms have the potential to play a significant role in shaping afferent input through these pathways. However, the relative role of ACh in regulating transmission in these pathways has not been examined. To compare cholinergic regulation of thalamic and cortical inputs, we injected an rAAV containing ChR2-EYFP under the control of the CaMKII promoter into the MTN of mice. After at least three weeks, brain slices were prepared and glutamatergic terminals in BLa from MTN and cortex were stimulated in the same slice and evoked field responses recorded at the same site. MTN fEPSPs were evoked by optogenetic stimulation of MTN terminals in BLa with single pulses of blue light, while cortical fEPSPs were evoked by electrical stimulation of cortical afferents in the external capsule (Fig 3E). The effect of tonic elevation of ACh was assessed in each pathway following application of physostigmine (10 μM). As previously observed (Fig. 4A), elevated tonic ACh strongly suppressed the cortical fEPSP (Fig. 4B). This inhibition was blocked by atropine (5 μM), indicating that it was mediated by muscarinic receptors. Subsequent application of mecamylamine (10 μM) had no additional effect, suggesting that nicotinic receptors were not involved. In contrast, at the same recording site elevation of tonic ACh with physostigmine had no significant effect on baseline responses to MTN pathway stimulation. However, application of atropine significantly increased the MTN fEPSP and this increase was blocked by mecamylamine. These findings suggest that at baseline, tonic ACh engaged both muscarine and nicotinic receptors to produce opposing and offsetting effects on the fEPSP. Application of atropine blocked the muscarinic inhibition revealing the unopposed nicotinic facilitation which was subsequently blocked by mecamylamine. These findings reveal distinct effects of muscarinic and nicotinic receptors in these pathways. During tonic ACh, cortical input was strongly inhibited by muscarinic receptors, but little affected by nicotinic receptors. In contrast, thalamic input was more strongly facilitated by nicotinic receptors with markedly less muscarinic inhibition than at cortical inputs.

### Differential regulation of cortical and thalamic input to the BLa by muscarinic receptors

To better examine pathway specific differences in the effect of ACh, we injected an rAAV containing ChR2-EYFP under the control of the CaMKII promoter into either the PL or the MTN of mice. After 3-4 weeks, brain slices were prepared and the effect of muscarine, a selective mAChR agonist, on fEPSPs evoked by a single blue light pulse to either PL or MTN terminals in the BLa examined. Muscarine (10 μM) inhibited fEPSPs in both pathways with no sex-dependent difference at either PL or MTN input (% Control; PL Males 18.2 + 2.2% (n = 26, N = 26); Female 20.8 + 4.2% (n = 6, N = 6), p = 0.6, student’s t-test; MTN Males 51.8 + 6.4% (n = 14, N = 14), Female 39.2 + 4.5% (n = 13, N = 13), p = 0.13, student’s t-test) so data were collapsed across males and females for all experiments. Increasing concentrations of muscarine (0.03 – 30 μM) produced a monotonic decrease in the amplitude of the fEPSP at both inputs (Fig. 3A). The effect of muscarine on PL-evoked fEPSPs could be fit to a standard logistic equation yielding an IC_50_ of 0.56 μM (95% c.i. 0.38 to 0.80 μM) and Hill coefficient of −0.58 (95% c.i. −0.68 to −0.48; n = 5-35 slices). Similar analysis of the MTN-evoked fEPSP indicated that the effect of muscarine in this pathway was shifted approximately 10 fold to the right (IC_50_ = 6.04 (95% c.i. 3.66 to 9.23 μM), Hill coefficient = 0.56 (95% c.i. −0.79 to −0.40; n = 4-27). The confidence intervals of the IC_50_ concentrations at these two inputs did not overlap indicating that PL input was significantly more sensitive to inhibition by muscarine than was MTN input.

Input to BLa from ventral subiculum (vSub) also plays an important role in regulating amygdalar responses to emotionally arousing stimuli. Given the differing effects of muscarine at PL and MTN inputs, we also assessed muscarine inhibition of input from vSub. fEPSPs were evoked by optogenetic stimulation of vSub terminals in BLa 4 weeks after injection into vSub of AAV containing ChR2-EYFP. Muscarine (10 μM) strongly inhibited these fEPSPs, similar to its effect on PL-evoked fEPSPs, but significantly greater than its inhibition of MTN inputs. As observed following EC stimulation (Fig 2F), the effect of muscarine on both PL and MTN inputs was presynaptic, since muscarine significantly enhanced the paired pulse ratio in both pathways (Fig 3C,D). Taken together, these results indicate the presynaptic nature of muscarinic inhibition and that PL and vSUB input to BLa are significantly more sensitive to this inhibition than MTN input.

### Acetylcholine acts through M3 and M4 receptors to suppress transmission

To identify the mAChR subtype(s) involved in the muscarine-mediated inhibition of the PL- and MTN-evoked fEPSP we used a protocol in which 10 min of baseline recording was followed by perfusion with muscarine (10 μM) to inhibit the fEPSP before addition of selective muscarinic receptor antagonists. Each drug was perfused until a steady state effect was observed before moving to the next drug. PL or MTN inputs were optogenetically stimulated and AMPA receptor fEPSPs were isolated using 10 μM picrotoxin, 2 μM CGP55845 and 50 μM APV or 10 μM MK-801 to block GABA and NMDA receptors. M1 receptors are the most abundant mAChR in the BLa and have been reported to be present at presynaptic glutamatergic terminals (Muller et al., 2013). However, at both the PL and MTN inputs the selective M1 receptor antagonist, telenzepine (100 nM) had no significant effect on the fEPSP in the presence of muscarine. Similarly, VU0255035 (5 μM), a selective M1 receptor antagonist with greater than 75-fold selectivity over M2–M5 receptors (Sheffler et al., 2009) also did not produce significant reversal of muscarinic inhibition in either pathway. Consequently, the effect of these two antagonists was combined (Figure 6A, B) and indicated little functional involvement of M1 receptors in the muscarinic inhibition. Similarly, M2 receptors were also not involved as the highly selective M2 receptor antagonist AF-DX 116 (1μM) had no effect on muscarinic inhibition in either pathway (Figure 6A, B).

**Figure 6.**
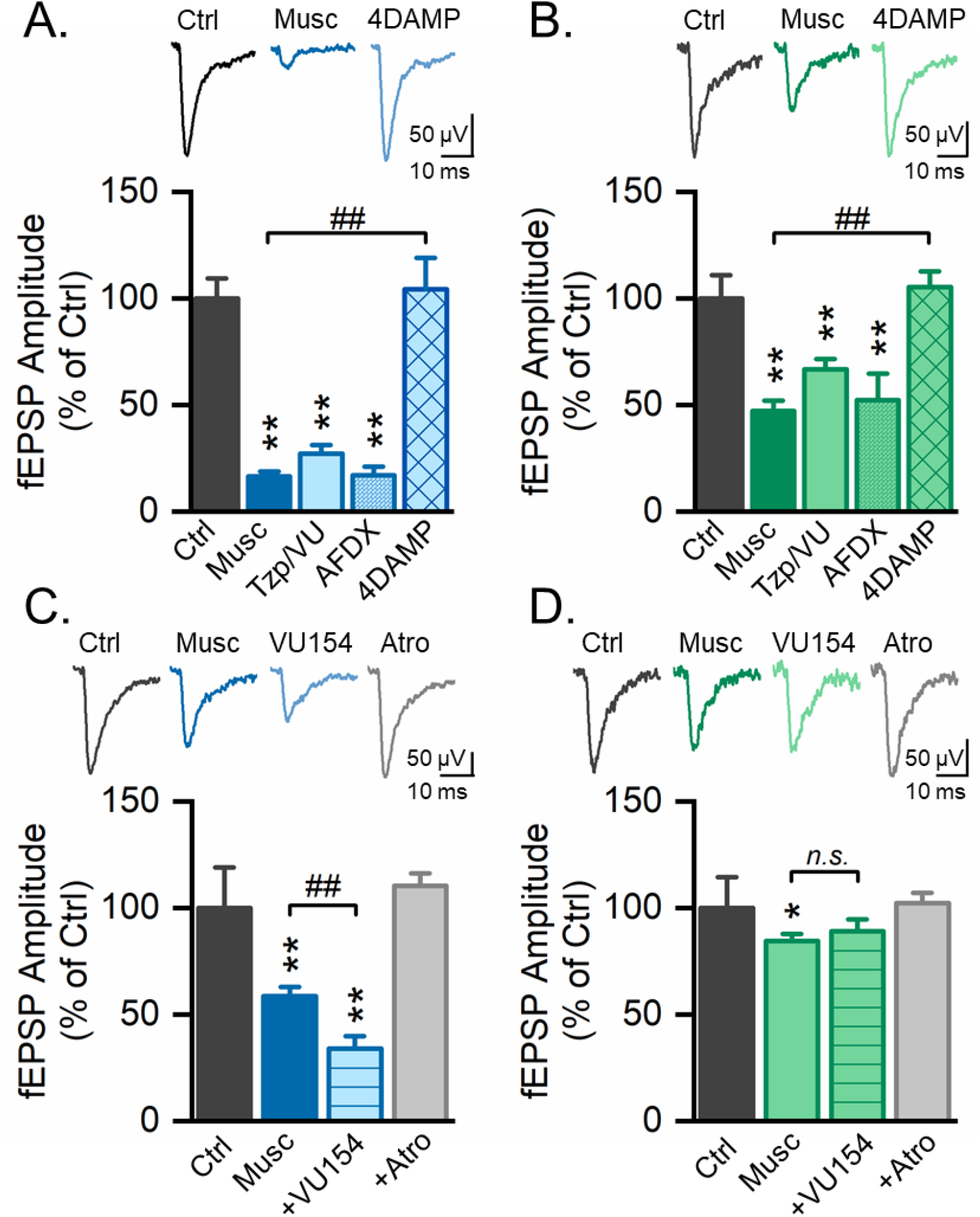
M3 and M4 mAChRs differentially regulate glutamatergic synaptic transmission from PL and MTN o the BLa**. A.** Antagonism of M1 receptors with telenzepine (Tzp, 100 nM, 27.8 + 6.5%, n = 6, N = 6) or VU0255035 (VU035, 5 μM, 25.8 + 1.4%, n = 3, N = 3) failed to reduce muscarine (10 μM) inhibition of PL input. Given the similarity in the lack of effect of these antagonists (Tzp 27.8 + 6.5% of baseline; VU035 25.8 + 1.4% of baseline), the results were combined. Antagonism of M2 receptors with AF-DX 116 (1 μM) also failed to reverse muscarinic inhibition of the fEPSP (n=6; N=6). In contrast, the M3/M4 antagonist 4DAMP (1μM) blocked muscarinic inhibition (n=10; N=10; One-way ANOVA, F(5,94)=45.90; p < 0.0001), indicating that M3 or M4 receptors were responsible for inhibition in the PL pathway. **B.** At MTN input muscarine (10 μM) produced less inhibition than at PL input. However, as at PL input, this inhibition was not reduced by M1 antagonists (TZP or VU035, n = 6, N = 6) or the M2 antagonist AF-DX 116 (n = 5, N = 5), but was reversed by 4-DAMP (n = 9, N = 9; One-way ANOVA, F(5,74)=31.48; p < 0.0001), indicating that M3 or M4 receptors were responsible. **C.** The M4 PAM, VU0467154 significantly potentiated inhibition produced by 0.3 μM muscarine in the PL pathway (n = 8, N = 8; One-way ANOVA, F(3,28)=59.86; p < 0.001), indicating a role for presynaptic M4 receptors. **D.** In contrast, at MTN input, muscarine (0.3 μM) produce a small but significant inhibition (n=4, N=4, One-way ANOVA, F(3,12)=4.62; p =0.023) and the M4 PAM did not potentiate this inhibition (p=0.85, post-hoc Tukey test), indicating that muscarine produced inhibition in this pathway by acting on M3 receptors. *p<0.05; **p<0.01, ##p<0.01, n.s. not significant.

In contrast, the M3 antagonist 4-DAMP (1 μM) completely reversed muscarinic inhibition at both pathways to the BLa (Fig. 6A,B). While 4-DAMP is considered an M3 antagonist, it shows limited selectivity over M1, M4, and M5 receptors (Dorje et al., 1991; Moriya et al., 1999; Watson et al., 1999). However, the inability of selective M1 or M2 receptor antagonists to block muscarinic inhibition and the lack of evidence supporting M5 receptors in the BLa (Lebois et al., 2018), suggests that 4-DAMP must block muscarinic inhibition by acting on either M3 or M4 receptors.

To investigate any contribution of M4 receptors to inhibition at PL and / or MTN input we used the highly selective M4 positive allosteric modulator (M4 PAM) VU0467154 (VU154). In these experiments a low-dose of muscarine (0.3 μM) was initially applied followed by VU154 (3 μM). At the PL pathway inhibition by this low dose of muscarine was significantly enhanced after application of the M4 PAM (Fig. 6C), indicating that presynaptic M4 receptors are present on PL terminals and inhibit glutamatergic transmission in this pathway. VU0154 (3 μM) also facilitated inhibition produced by another muscarinic agonist, oxotremorine. The M4 PAM increased oxotremorine (0.3μM)-induced inhibition from 18.7 ± 2.8% in baseline to 60.4 ± 8.6% in the presence of the M4 PAM (n=5; N=5; p=0.013, paired t-test). In contrast, the M4 PAM had no effect on either muscarine-induced (Figure 6D) or oxotremorine (5 μM)-induced inhibition (18.9 + 2.5% inhibition in oxotremorine, 18.2 + 1.1% inhibition in oxotremorine+VU154; n=3; N=3; p = 0.76, paired t-test) at the MTN input. Taken together, these experiments suggest that M4 receptors contribute to muscarinic inhibition at PL input to BLa, while inhibition at MTN inputs is exclusively mediated by M3 receptors.

### Muscarine inhibits synaptic transmission in the PL pathway through Gi/o protein-coupled M4 mAChRs

Because M3 receptors couple to Gq proteins and M4 receptors to Gi/o proteins, treating slices with an agent that inhibits Gi/o proteins should distinguish between inhibitory effects mediated by M3 and M4 receptors. Therefore, to further confirm a role for M4 receptors in producing inhibition in the PL pathway, we assessed the effect of Gi/o protein inactivation by bath application of the sulfhydryl alkylating agent n-ethylmaleimide (NEM) on the effects of muscarine (Shapiro et al., 1994; Morishita et al., 1997). Baclofen, a GABA_B_ receptor agonist that inhibits glutamate release through a Gi/o coupled mechanism in the BLa (Yamada et al., 1999) served as a positive control. As expected, baclofen (10 μM) significantly inhibited the fEPSP evoked by optogenetic stimulation of the PL input and this inhibition was reversed by the selective GABA_B_ antagonist, CGP55845 (2 μM, Fig. 7A). In separate experiments we then used a protocol in which 10 min of baseline recording was followed by perfusion with muscarine (10 μM) to inhibit the PL-evoked fEPSP and establish the baseline level of muscarinic inhibition. Muscarine was then washed out and NEM (50μM, Shapiro et al., 1994) was bath applied to slices for a minimum of 15 minutes. Muscarine (10μM) was again applied and the amplitude of the fEPSP after NEM treatment was compared to the fEPSP amplitude before NEM treatment. Baclofen (10 μM) was also applied following NEM treatment as a positive control and the extent of inhibition compared to that produced by baclofen in the absence of NEM in additional brain slices from the same animals. Incubation of slices with NEM was sufficient to inactivate Gi/o proteins, as effects of baclofen were significantly inhibited (Fig. 7B). Similar to its effects on baclofen inhibition, NEM also blocked muscarine inhibition (Fig. 7B). The similarity in the effect of NEM on baclofen and muscarine inhibition suggests that both agents act at PL input through Gi/o protein dependent mechanisms and supports the conclusion that muscarine inhibits glutamate release at PL input through Gi/o coupled presynaptic M4 receptors.

**Figure 7.**
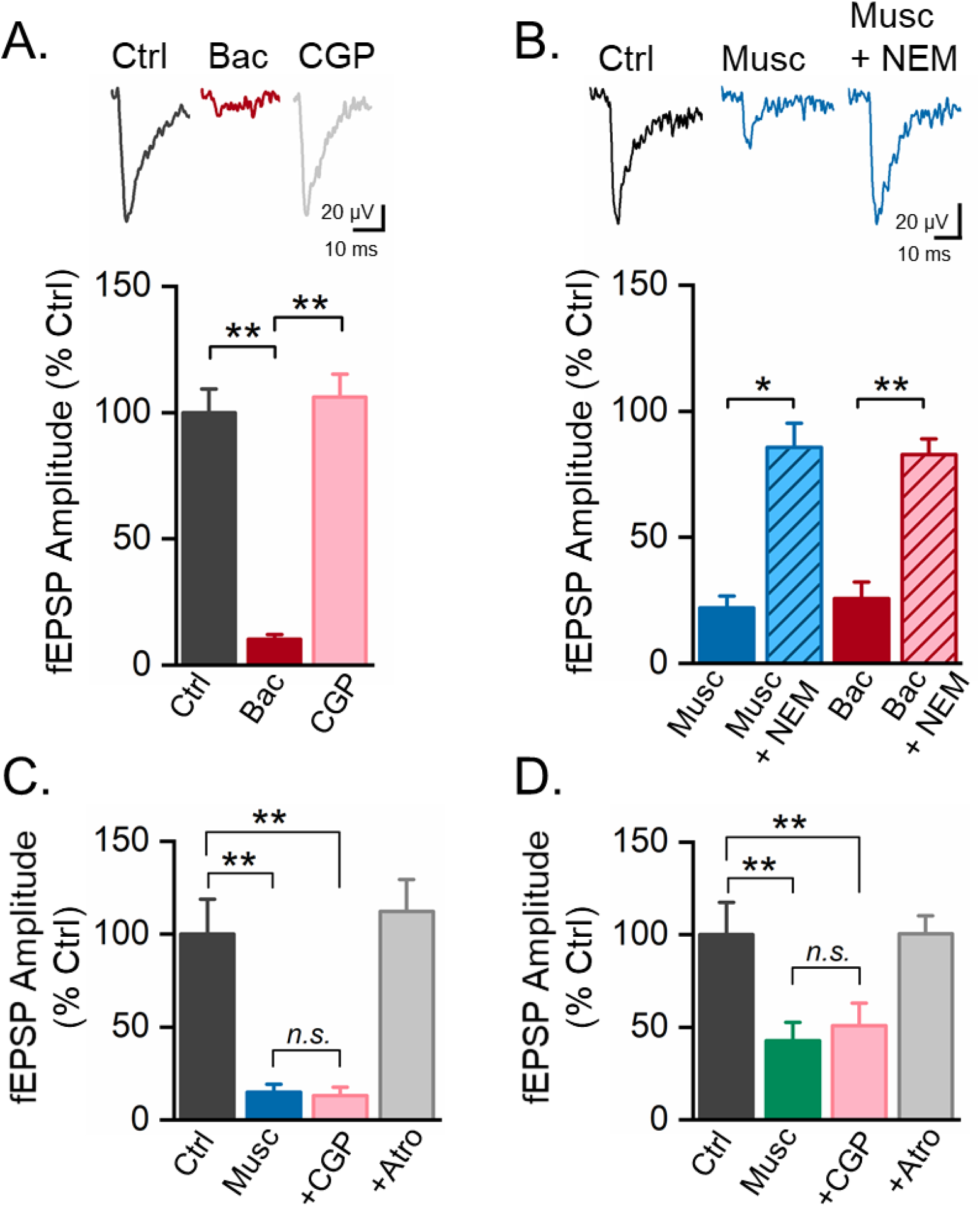
Mechanisms of muscarinic inhibition. **A, B.** Muscarine inhibition of PL input is dependent upon Gi/o protein dependent signaling. **A.** Baclofen (10 μM), which acts through GABA_B_ receptors coupled to Gi/o proteins, inhibits fEPSPs evoked by optogenetic stimulation of the PL pathway. This inhibition is reversed by the GABAB receptor antagonist, CGP 55845 (CGP, 2 μM, n = 7, N = 7; One-way ANOVA, F(2,18)=102.495; p < 0.0001). **B.** Treatment of brain slices with n-ethylmaleimide (NEM, 50 μM) for 15 minutes inactivated Gi/o proteins and blocked muscarine (10 μM) inhibition in the PL pathway (n = 4, N = 4). In the same slices NEM also blocked the inhibitory effect of baclofen (10 μM) in this pathway, demonstrating that Gi/o proteins were inactive (n = 4, N = 4; One way ANOVA, F(4,15)=37.64; p < 0.0001). These findings suggest that muscarine inhibition in the PL pathway is dependent upon Gi/o protein-coupled M4 receptors, rather than Gq protein-coupled M3 receptors. **C, D.** An alternative interpretation of these data is that muscarine produces inhibition indirectly through an M3 muscarinic receptor-mediated increase in inhibitory interneuron excitability, which activates GABAB receptors to suppress synaptic transmission. NEM would then block muscarine inhibition by blocking GABA_B_ receptor signaling. However, application of CGP55845 (2 μM) did not block muscarine inhibition in either the PL (2 μM, n = 7, N = 7; One-way ANOVA, F(3,24)=34.7; p < 0.0001) or MTN pathway (2 μM, n = 5, N = 5; One-way ANOVA, F(3,16)=11.39; p < 0.001). These findings indicate that muscarinic inhibition of PL and MTN inputs to BLa is not dependent on GABA_B_ receptors. *p<0.05, **p<0.01., n.s. not significant.

### M3 muscarinic receptors produce inhibition in the PL and MTN pathway through a mechanism that is independent of GABA_B_ receptors

An alternative explanation for the above findings is that muscarine acts on M3 receptors on GABAergic interneurons to increase interneuron excitability, releasing GABA which acts on GABA_B_ receptors to suppress synaptic transmission. This mechanism has recently been reported in hippocampal area CA1 (Goswamee and McQuiston, 2019). NEM would suppress this effect by blocking the action of Gi/o protein-coupled GABA_B_ receptors. However, as our experiments are performed in the presence of picrotoxin and CGP 55845, GABAA and GABAB receptors were not required for muscarinic inhibition at PL or MTN inputs to BLa. To determine if muscarinic inhibition was greater when GABAB receptors were available, we compared the extent of inhibition by muscarine in the absence and presence of CGP55845. Bath application of CGP55845 (2 μM) had no effect on muscarine inhibition in either pathway, indicating that even though presynaptic GABAB receptors are present, muscarine suppression of glutamatergic fEPSPs at PL and MTN inputs is independent of GABA_B_ receptors (Fig. 7C, D). Similarly, in whole cell experiments blockade of GABAergic inhibition by addition of picrotoxin (50 μM) and CGP 55845 (5 μM) did not alter either the early facilitation (ACSF, 111.3 + 2.8% vs GABA blockers, 110.7 + 2.0%, n = 3, p = 0.87, paired t-test) or late inhibition (ACSF, 84.5 + 3.3% vs GABA blockers, 82.2 + 3.7%, n = 5, p = 0.2, paired t-test) produced by stimulation of cholinergic terminals, indicating that acetylcholine did not act through GABAergic mechanisms to produce its effects.

### Muscarine inhibits MTN inputs through an M3 receptor dependent facilitation of retrograde endocannabinoid signaling

Endocannabinoids (eCBs) serve a retrograde inhibitory role in many brain regions (Ohno-Shosaku and Kano, 2014), allowing neurons to regulate their upstream neuronal inputs. Postsynaptic Gq-coupled muscarinic (M1/M3) receptors can facilitate retrograde eCB release, suppressing GABA (Kim et al., 2002; Ohno-Shosaku et al., 2003) or glutamate transmission (Chiu and Castillo, 2008; Kodirov et al., 2009). While this mechanism has not previously been reported at excitatory terminals in the BLa, it is possible that postsynaptic M3 receptors on BLa pyramidal cells could act through retrograde eCB release to inhibit glutamatergic transmission in the MTN or PL pathway. To examine this possibility, the selective CB1 antagonist, AM251 (1 μM) was applied in the presence of muscarine. At PL input, antagonism of CB1 receptors had no effect on muscarine inhibition (Fig. 8A). This lack of effect was somewhat surprising given the presence of CB1 receptors at these inputs, as application of CB1 receptor agonist WIN55,212 (5 μM) suppressed PL-evoked fEPSPs in a manner reversible by AM251 (Fig. 8B). These data suggest that even though CB1 receptors can inhibit PL evoked fEPSPs in the BLa, muscarinic suppression of PL input is not CB1 receptor dependent. Given the presence of CB2 receptors in the brain (Onaivi et al., 2008) and the ability of CB2 receptors to suppress transmitter release in some brain regions (Foster et al., 2016), in separate experiments we also tested the effect of the CB2 antagonist, AM630. However, as with CB1 antagonists, AM630 (2 μM) had no effect on muscarine inhibition (Musc: 30.1 + 5.2%; Musc+AM630: 27.7+2.0%; n=3; N = 3; p=0.75, paired t-test). In contrast, at MTN inputs blockade of CB1 receptors with AM251 completely reversed muscarinic inhibition of fEPSPs (Fig. 8C), while having no effect on baseline fEPSPs in the absence of muscarine (Fig. 8D). Muscarine inhibition at MTN input is dependent upon M3 receptors (Fig. 6). These findings suggest that at MTN inputs, muscarine inhibition is mediated by a postsynaptic M3 receptor-mediated release of eCBs which retrogradely acts on CB1 receptors on MTN terminals to inhibit glutamatergic transmission.

**Figure 8.**
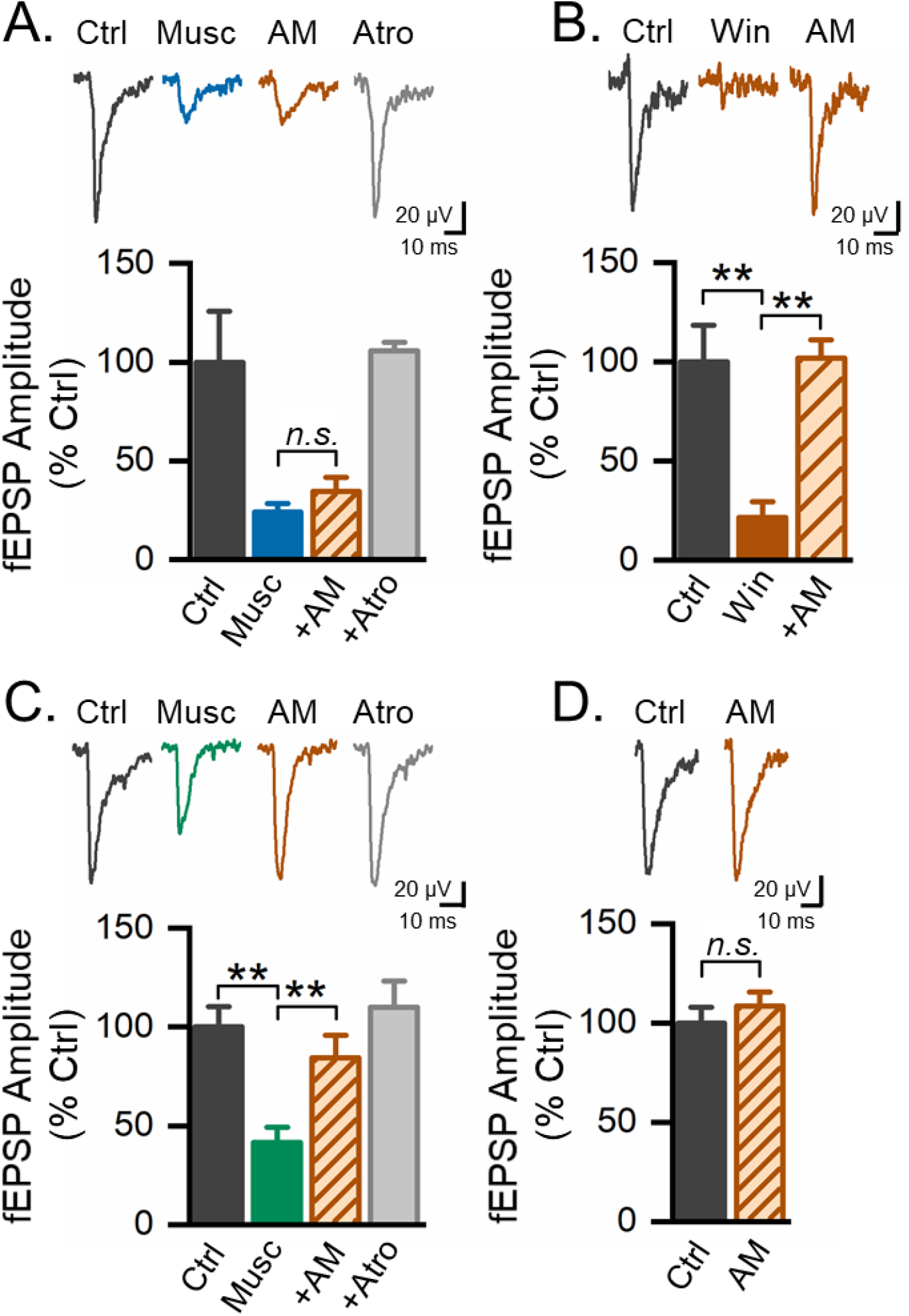
Muscarinic inhibition of MTN, but not PL input, is mediated by an M3 receptor dependent facilitation of retrograde endocannabinoid signaling. **A.** The CB1 receptor antagonist, AM251 (AM, 1 μM) had no significant effect on muscarine (10 μM) inhibition in the PL pathway. Muscarine inhibition was completely reversed by atropine (5 μM, n=7, N=7; One-way ANOVA, F(3,24)=84.90; p < 0.0001). **B.** The CB1 agonist, Win 55212-2 (5 μM) strongly suppressed fEPSPs at PL inputs. This suppression was reversed by AM251, indicating that it was dependent on CB1 receptors (n=6, N=6; One-way ANOVA, F(2,13)=53.03; p < 0.0001). These findings suggest that CB1 receptors are present at PL terminals, but are not engaged during muscarinic inhibition. **C.** In contrast, at MTN input AM251 (1 μM) reversed muscarine inhibition (n=7, N=7; One-way ANOVA, F(3,24)=9.90; p < 0.0002). Subsequent addition of atropine had no additional significant effect (Tukey post-hoc test, p=0.26). **D.** AM251 by itself had no significant effect on the optogenetically-evoked fEPSP at the MTN input (n=3, N=3, p=0.34, paired t-test), indicating that AM251 did not directly facilitate synaptic transmission in this pathway. Together, these results suggest that muscarine inhibits responses in the MTN pathway by acting on postsynaptic M3 receptors to facilitate retrograde endocannabinoid release which acts on presynaptic CB1 receptors on MTN terminals. **p<0.01., n.s. not significant.

### Frequency-dependent inhibition of glutamatergic input by mAChRs

PL and MTN inputs are differentially modulated by mAChRs in response to single pulse stimulation. However, theta (4-12 Hz) and gamma (30-80 Hz) frequency activity occur in the BLa during emotional behavior and associative learning (Stujenske et al., 2014; Bocchio et al., 2017), making it of considerable interest to understand how ACh regulates afferent synaptic transmission at different frequencies in each pathway. Therefore, we investigated the effect of muscarine on responses in PL and MTN pathways to short stimulus trains at frequencies within a behaviorally relevant range *in vivo.* Stimulus trains consisting of 10 light pulses were delivered to either input at frequencies ranging from 1-40 Hz in the absence or presence of muscarine (10 μM). At PL synapses, stimulation at 1 Hz evoked responses of similar amplitude throughout the train. Muscarine (10 μM) strongly and similarly suppressed each response of the train (Fig., 9A, B). Alternately, stimulation at 40 Hz evoked a facilitation on the second response of the train (Fig 9A, B) in line with earlier results showing paired pulse facilitation in this pathway (Figs 3 & 5). Subsequent pulses in the train evoked progressively smaller fEPSPs such that the last fEPSP was 35.5 + 2.8% of the amplitude of the first fEPSP. Following addition of muscarine, the first response of the train was strongly inhibited, as seen with single pulses, but subsequent responses were facilitated relative to the first fEPSP. This facilitation was maintained throughout the remainder of the train, such that the fEPSP amplitude in response to the last pulse of the train in muscarine was similar to the fEPSP amplitude to the last pulse in control (Fig. 9A, B), reflecting a complete loss of muscarine inhibition during the train. When comparing the extent of muscarine inhibition on the last pulse of different frequency trains, it could be seen that muscarine inhibition during the train was frequency dependent (Fig. 9C). Inhibition was preserved during low frequency 1 Hz stimulation, but increasingly attenuated as the frequency of the train increased. At 40 Hz, a frequency in the gamma range, inhibition was completely lost during the train. A similar result was also found at MTN input. As seen with single pulses, muscarine inhibition was significantly less in this pathway compared to PL input. However, as in the PL pathway, this muscarine inhibition was preserved during low frequency (1-5 Hz) trains, but attenuated during trains with frequencies greater than 5 Hz, reaching a complete loss of inhibition at 40 Hz. Thus, at both PL and MTN inputs, muscarinic receptors act as a high pass filter, blocking low frequency signals, while allowing higher frequency signals to reach the BLa.

**Figure 9.**
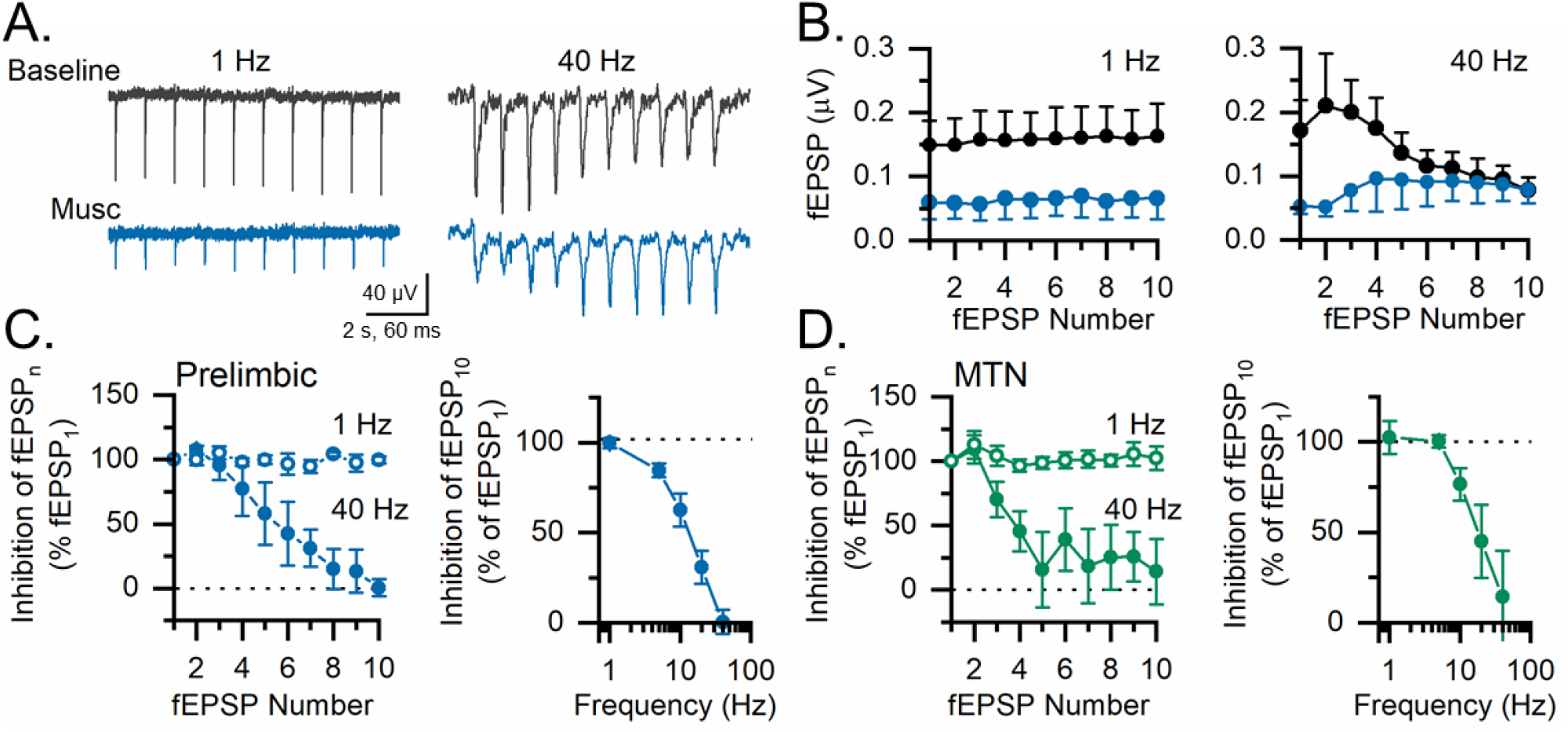
Muscarine inhibition at PL and MTN inputs is frequency dependent. **A, B.** Optogenetic stimulation of PL input at 1 Hz evoked fEPSPs that were consistently inhibited by muscarine (10 μM) during the stimulus train. During 40 Hz stimulation, fEPSPs in baseline initially facilitated then became progressively smaller. In muscarine fEPSPs were initially strongly inhibited. They then facilitated for the remainder of the train such that by the end of the train the fEPSP in muscarine was the same amplitude as in control, reflecting a loss of muscarine inhibition (n=4, N=4). **C.** At low frequency (1 Hz) muscarine inhibition was stable during the train. At high frequency (40 Hz) muscarine inhibition progressively declined with subsequent pulses until it was completely absent by the end of the train. Muscarine inhibition at the end of the 10 pulse stimulus train progressively declined at higher stimulus frequencies. **D.** A similar frequency dependence of muscarine inhibition was also observed at MTN input (n = 6, N=6).

## Discussion

Our results show robust ACh regulation of afferent input to BLa that depends upon the mode of ACh release, and is pathway specific and frequency dependent. Single pulse stimulation of cholinergic terminals engaged both nAChRs and mAChRs, producing a biphasic excitatory-inhibitory modulation of cortical input in the BLa. By contrast, elevation of extracellular ACh by blockade of acetylcholinesterase produced solely monophasic muscarinic inhibition of cortical input. At thalamic input, this same increase in extracellular ACh had no net effect on synaptic transmission. The differences in sensitivity of cortical and thalamic inputs to muscarinic inhibition were attributed to distinct mechanisms of mAChR action at each site. Muscarine inhibition at both inputs disappeared at higher frequencies of stimulation, consistent with its action as a high pass filter for afferent BLa signals.

Pharmacological studies with persistent agonist application have demonstrated that both nicotinic and muscarinic receptors regulate transmitter release in the BLa (Sugita et al., 1991; Yajeya et al., 2000; Jiang and Role, 2008). The present study extends those findings by showing rapid regulation of glutamatergic transmission by endogenously released ACh. These findings are consistent with anatomical studies showing cholinergic terminals converging on glutamatergic synapses in BLa (Li et al., 2001; Muller et al., 2011). Single pulse stimulation of cholinergic terminals produced an immediate (<20 ms) and short-lived nAChR-mediated facilitation of cortical input to BLa, followed by a slower mAChR-mediated inhibition, lasting for up to 1 sec. Both facilitation and inhibition of afferent input were evoked by the same single cholinergic stimulus. Prior studies have reported postsynaptic responses to individual cholinergic stimuli in inhibitory neurons in thalamus and cortex (Sun et al., 2013; Urban-Ciecko et al., 2018). However, to our knowledge this is the first study that demonstrates that ACh release can potentiate glutamate release on the timescale of an individual synaptic event. This is also the first demonstration of this form of excitatory-inhibitory neuromodulation by ACh in the amygdala and suggests that cholinergic neuromodulation can serve precise, computational roles in the BLa network. The presence of these forms of cholinergic modulation in BLa is consistent with the robust cholinergic innervation of this region and further supports the vital role of ACh in information processing in this region.

Cholinergic neurons in BF exhibit fast and precise responses to both appetitive and aversive behavioral cues (Hangya et al., 2015). Studies using fluorescent ACh sensors have found that during emotionally salient stimuli, there is a phasic release of acetylcholine into the BLa (Crouse et al., 2020) that can mediate associative learning (Jiang et al., 2016). In addition, phasic BF cholinergic stimulation can induce acute appetitive behaviors (Aitta-aho et al., 2018). It is tempting to speculate that the excitatory-inhibitory modulation of glutamatergic transmission by endogenously released ACh observed here reflects the action of phasically released ACh in the BLa during these behaviors. This phasic ACh rapidly engaged nAChRs on cortical terminals in BLa to facilitate glutamate release for up to 20 ms following cholinergic terminal activation. In contrast, glutamate release produced by action potentials arriving 50-1000 ms after simulation of cholinergic inputs was suppressed by robust mAChR-mediated inhibition. Together, the biphasic action of endogenously released ACh on presynaptic nicotinic and muscarinic receptors suggests that it would entrain glutamatergic input in a tight temporal window following cholinergic terminal activation and suppress poorly timed input that arrived outside of this window. This mechanism would enhance the signal-to-noise ratio for cortical input to BLa, thereby facilitating attention to salient signals (Bloem et al., 2014; Dannenberg et al., 2017) and may be important in forms of heterosynaptic plasticity in the BLa (Jiang et al., 2016).

Acetylcholine release from the BF occurs at multiple physiological timescales, ranging from milliseconds to minutes and hours (Disney and Higley, 2020; Sarter and Lustig, 2020). To better understand the consequences of elevated tonic acetylcholine on glutamate transmission, we increased extracellular ACh by inhibiting acetylcholinesterase with physostigmine. In contrast to phasic ACh, tonic ACh evoked a steady state and reversible monophasic inhibition of cortical input mediated entirely by mAChRs. The lack of nAChR involvement is likely attributed to nAChR desensitization during sustained ACh, which has been well documented for these receptors (Quick and Lester, 2002; Giniatullin et al., 2005). These findings suggest that during behavioral states associated with high cholinergic tone, ACh regulation of cortical input would be predominantly inhibitory. In contrast, tonic ACh produced little net effect at MTN input. The lack of effect was associated with both a larger persistent nicotinic facilitation and a smaller muscarinic inhibition that opposed and occluded each other. The persistence of nAChR mediated facilitation at MTN input during elevated tonic ACh differs from that at PL inputs. This difference may reflect distinct nAChR receptor types at MTN compared to PL synapses (Quick and Lester, 2002; Venkatesan and Lambe, 2020) or differences in the anatomical arrangement of cholinergic release sites and thalamic terminals (Disney and Higley, 2020). This could result in lower concentrations of ACh at MTN synapses which would be less likely to desensitize nAChRs.

In addition to differences in nicotinic facilitation, MTN synapses were also subject to significantly less muscarinic inhibition than PL input. This disparity was caused by differential regulation of transmitter release by M4 and M3 receptors at the two inputs. These findings are consistent with growing evidence of highly specific localization of muscarinic receptor types to distinct neural pathways in the brain (Gil et al., 1997; Palacios-Filardo et al., 2021). The finding that M4 receptors regulate PL input is the first demonstration of presynaptic inhibition by M4 receptors in the BLa and builds on prior work showing presynaptic regulation by M4 receptors in other brain regions (Dasari and Gulledge, 2011; Pancani et al., 2014; Yang et al., 2020; Palacios-Filardo et al., 2021). Inhibition by M4 receptors was likely mediated by a suppression of presynaptic N- and P-type voltage-gated calcium channels through a Gi/o protein-dependent mechanism (Hille, 1994; Howe and Surmeier, 1995; Yan and Surmeier, 1996). Blockade of muscarinic inhibition by NEM in the present study supports this conclusion (see also Shapiro et al., 1994). Muscarinic modulation of these calcium channels is voltage dependent and is attenuated by membrane depolarization (Yan and Surmeier, 1996). This voltage dependence could underlie the observed loss of muscarinic inhibition during high frequency stimulation when the presynaptic membrane would be depolarized. This mechanism could explain why low frequency transmission at cortical inputs would be suppressed by presynaptic mAChRs, but high frequency or burst transmission would pass. Presynaptic mAChRs would thereby serve as a high pass filter for incoming salient information from cortex.

In contrast, at MTN inputs muscarinic inhibition is mediated by M3 receptors. The differences in muscarinic receptor type at PL and MTN inputs provides a mechanism for differential sensitivity to ACh in these two pathways and is consistent with the finding that muscarine was significantly less potent at MTN than PL input. Our data indicate that ACh suppressed glutamate release at MTN inputs by acting on postsynaptic M3 receptors to stimulate retrograde eCB release which subsequently engaged CB1 receptors on thalamic terminals. This conclusion is supported by the ability of an M3 receptor antagonist to block the inhibition and the inability of muscarine to produce inhibition in the presence of a CB1 receptor antagonist. Muscarinic receptor-induced suppression of excitation (MSE) has been reported (Chiu and Castillo, 2008; Kodirov et al., 2009), but has not previously been demonstrated in BLa. However, its role in this region is consistent with both the high expression of CB1 receptors in the amygdala (Marsicano and Lutz, 1999) and the presence of these receptors in glutamatergic terminals in this area (Domenici et al., 2006; Fitzgerald et al., 2019). Our data show that a CB1 receptor agonist suppressed transmission at both PL and MTN inputs, indicating the presence of CB1 receptors at both glutamatergic synapses. The presence of MSE only at MTN input thus reflects the localization of M3 receptors capable of stimulating eCB release. These findings highlight the pathway specific control of glutamate release by distinct cholinergic receptors and provide targets to selectively modulate individual components of acetylcholine’s actions.

The marked difference in the ability of tonic ACh to suppress transmission at PL and MTN inputs suggests that during behavioral states associated with high cholinergic tone, thalamic input will more strongly influence BLa activity than will cortical input. These findings are consistent with prior work in cortex showing that ACh enhances the influence of thalamic sensory input on cortical activity through a nicotinic facilitation of glutamate release, and reduces internal corticocortical connections by presynaptic muscarinic inhibition (Hasselmo, 2006; Hasselmo and Sarter, 2011). The resulting reduction in cortical feedback excitation is postulated to reduce interference from previous retrieval and thereby enhance memory encoding and attention to novel sensory input. The differential nicotinic and muscarinic modulation of PL and MTN inputs seen in the present study may similarly favor thalamic sensory input and reduce cortical feedback in amygdala during behavioral states associated with high cholinergic tone. In this way ACh would prioritize amygdala inputs to facilitate encoding of emotional memories and attention to novel cues.

## Acknowledgments

This work was supported by the NIMH (R01MH104638 to DDM and AJM), an ASPIRE2 Award from UofSC VP for Research (DDM), a Support to Promote the Advancement of Research and Creativity (SPARC) Graduate Research Grant from the UofSC VP for Research, Grant Number T32-GM081740 from the NIH-NIGMS (SCT) and partly supported by Merit Award # I01 BX001374 to Marlene A. Wilson (PI) from the United States Department of Veterans Affairs Biomedical Laboratory Research and Development Service (VA BLRD).

